# Spatial Transcriptomics Reveals Host Gene Expression Patterns Associated with *Naegleria fowleri* Infection in the Murine Brain

**DOI:** 10.64898/2026.11.18.767516

**Authors:** Joseph Dainis, Michael Bailey, Mackenzie A. C. Sievert, Anthony A. Ruberto, Dennis E. Kyle

## Abstract

Primary Amoebic Meningoencephalitis (PAM), caused by the free-living amoebae *Naegleria fowleri*, is a devastating disease with a mortality rate exceeding 97%. Despite its severity, the spatial and temporal characteristics of the host immune response to infection within the brain remain poorly understood. Here, we used spatially resolved transcriptomic profiling to characterize region-specific gene expression patterns in brain tissue collected at multiple time points following intranasal inoculation of *N. fowleri* in mice. Our analysis revealed pronounced pro-inflammatory transcriptional signatures that varied across brain regions and over the course of infection. Cell type-associated gene expression profiles suggested the involvement of several immune cell populations, including macrophages, neutrophils, and natural killer T cells, alongside transcriptional changes associated with neural cell populations. Notably, inflammatory transcriptional signatures also were detected in brain regions without detectable amoebae, indicating that host responses extend beyond regions of parasite localization. Collectively, these findings provide a spatial and temporal characterization of the host transcriptional response during experimental PAM, highlighting inflammatory gene expression signatures and putative immune and neural cell populations associated with disease progression. This study provides a foundation for future investigations into the cellular and molecular processes underlying neuroinflammation during *N. fowleri* infection.

**IMPORTANCE:** *Naegleria fowleri*, commonly known as the brain-eating amoeba, causes a rapidly progressing brain infection that is fatal in more than 97% of cases. In this study, we examined gene expression patterns across different brain regions and stages of infection in mice to better understand the host response. We identified expression signatures consistent with developing inflammation and predicted changes in the relative contributions of different immune cell populations. Notably, following invasion of the brain by *N. fowleri*, immune-associated gene expression signatures were also detected in regions without detectable amoebae, suggesting that host responses extend beyond areas of direct invasion. Together, these findings provide new insights into the spatial and temporal progression of host immune responses during *N. fowleri* infection and identify pathways for further investigation that may inform the development of host-directed treatments.

## INTRODUCTION

*Naegleria fowleri*, also known as the brain-eating amoeba, is a free-living eukaryotic pathogen that causes Primary Amoebic Meningoencephalitis (PAM). Humans are susceptible to infection when the trophozoite form of the parasite, found in warm freshwater globally, enters the nose, where the amoebae invade the olfactory epithelium, traverse the olfactory nerves, and enter the frontal lobe of the brain (1). There, amoebae feed on neural tissue, inducing a rare disease that results in mortality in >97% of cases (2). Thirty-nine countries have documented cases of PAM, with the United States, India, Pakistan, Australia, Mexico, and the Czech Republic reporting the majority of cases (3). In the United States, up to eight cases of PAM are reported each year, a number predicted to increase with global warming (4, 5).

A major contributor to the high mortality of confirmed PAM infections is the lack of effective clinical treatments, reflecting persistent gaps in our understanding of the host and parasite factors that shape disease (6, 7). Previous studies investigating host responses to *N. fowleri* infection have largely focused on innate immunity (8, 7). Among innate immune cell populations, neutrophils have been consistently identified as prominent components of the inflammatory response in the brains of experimentally infected animals and within brain lesions from human patients (9, 10). Neutrophil predominance has also been observed in the cerebrospinal fluid of infected patients (11). Furthermore, neutrophils exhibit anti-amoebic activity *in vitro*, suggesting a potential role in the host immune response to *N. fowleri* (12). Macrophages and brain-resident microglia also exhibit anti-*N. fowleri* activity *in vitro* through cytolytic mechanisms and the production of proinflammatory cytokines (13, 14, 15).

Although these studies have provided insights into the cellular components of the host immune response to *N. fowleri*, how these responses develop and are distributed within the brain remains poorly understood. Recent bulk transcriptomic analyses of brain tissue from mice infected with *N. fowleri* isolates of differing virulence revealed widespread changes in gene expression associated with immune responses and neural regulation (16). However, these analyses provide limited insight into the temporal progression of host responses and may obscure localized transcriptional differences between regions with and without detectable amoebae.

Advances in spatial transcriptomics offer opportunities to address these limitations by resolving gene expression within intact tissue architecture (17, 18). Applied across multiple time points, these approaches can characterize the spatial distribution and temporal progression of host responses during infection. Spatial transcriptomics has been used to investigate tissue-specific responses to diverse infectious agents (19), including Japanese encephalitis virus (20), trypanosomatids (21), and *Toxocara* spp. (22), as well as responses to adeno-associated viral vectors in the murine brain (23).

In this study, we investigated changes in host gene expression and associated pathways in *N. fowleri*-infected mouse brain tissue, with particular emphasis on their spatial distribution and temporal dynamics during disease progression. We characterized host transcriptional profiles across spatially defined regions with varying amoeba burdens at successive time points during infection. Together, these findings provide insight into the spatial and temporal development of host responses to *N. fowleri* infection and identify transcriptional signatures associated with the progression of PAM.

## RESULTS

### Spatial transcriptomic profiling and temporal distribution of amoebae in the mouse brain

We generated a spatial transcriptomic dataset from brain sections collected from ten mice inoculated with 5,000 *N. fowleri* V596 amoebae and two mock-infected mice across a four-day infection course (**Fig 1A**). We used antibody staining to visualize amoebae and microglial cells and to guide the selection of regions of interest (ROIs) for transcriptomic profiling (**Fig 1B**; **Fig S1**). We first classified ROIs by amoeba burden using tissue collected at day 4 post-infection (p.i.; N = 3 tissue sections; n = 24 ROIs), establishing three groups: no detectable amoebae, low amoeba burden (1–9 amoebae/ROI), and high amoeba burden (*≥* 10 amoebae/ROI) (**Fig 1C**).

**FIG 1.**
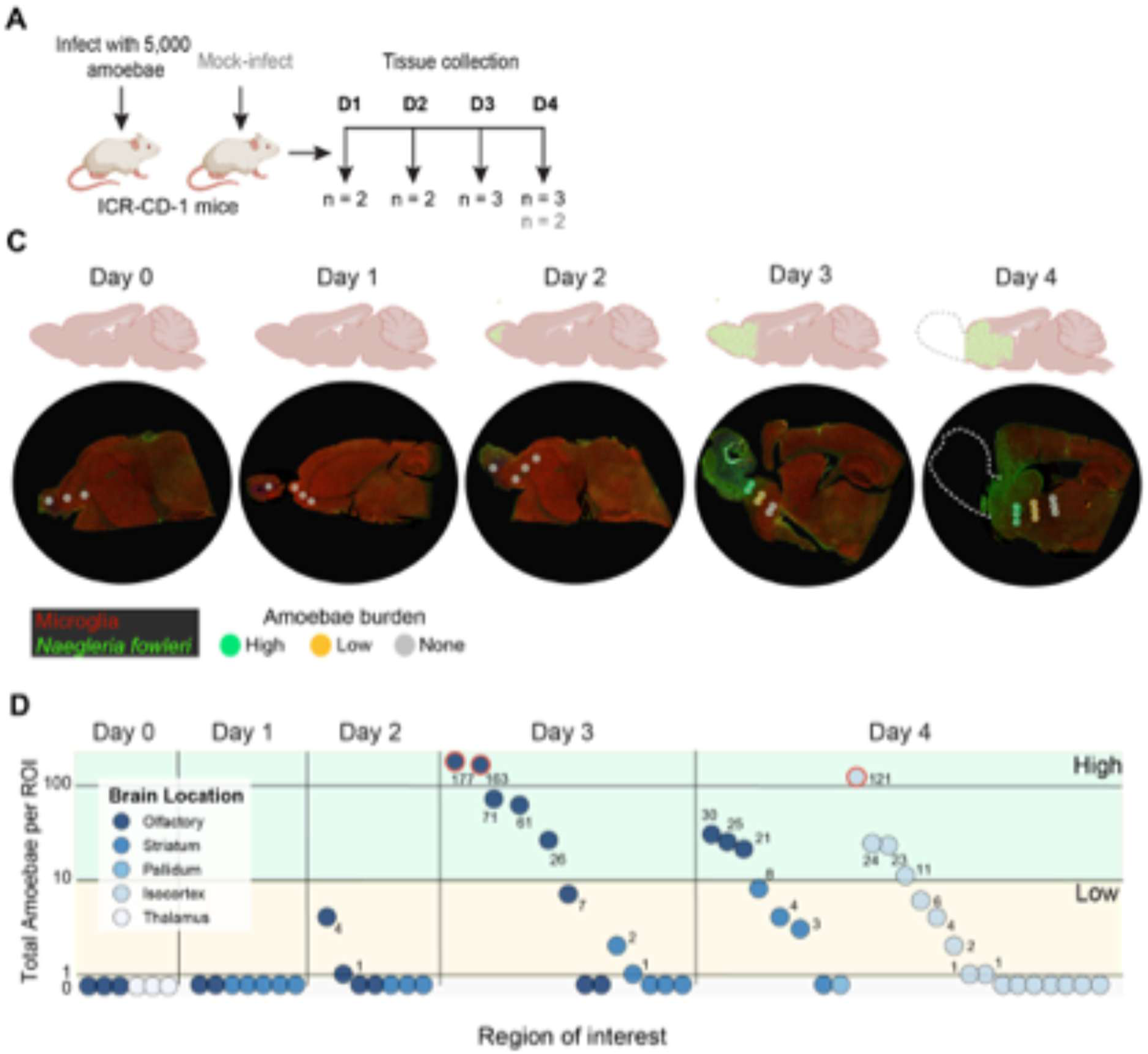
Quantification of amoeba burden in *N. fowleri*-infected mouse brains over time. (A) Schematic of the experimental design. Ten female CD-1/ICR mice were infected with 5,000 Vero cell-passaged N. fowleri V596 amoebae, while two mice were mock-infected. Two to three mice were euthanized daily throughout the course of infection, which typically progresses to end-stage disease within 4 days in this mouse background. Following euthanasia, brains were collected, fixed in 10% formalin, and paraffin-embedded for tissue sectioning. (B) Placement of regions of interest (ROIs) within brain tissue sections. One to three ROIs were selected from immunostained brain sections based on estimated amoeba burden. ROIs were anatomically matched across tissue sections from different animals to account for regional differences in neuronal gene expression.(C) Representative images of uninfected and N. fowleri-infected brain tissue. Brain sections from 12 mice were fluorescently stained to visualize amoebae (green), microglia (red), and neutrophils (blue). (D) Quantification of amoebae within individual ROIs grouped by anatomical brain region. Brain regions were assigned using the Allen Mouse Brain Atlas as a reference (Figure S3A). ROIs circled in red were classified as neutrophil ROIs, as the transcripts isolated from these ROIs were only taken from areas of the ROIs with neutrophil staining, and were excluded from further analysis.

We then selected anatomically comparable ROIs from earlier infection time points and mock-infected tissue. Gene expression was quantified within each ROI using GeoMx Digital Spatial Profiler software (**Materials and methods**). In total, we profiled 57 ROIs, including 54 whole-area ROIs and three ROIs enriched for neutrophil-stained regions. Sequencing depth and transcript diversity for each ROI are summarized in **Table S1**.

To characterize the spatial and temporal distribution of amoebae during infection, we quantified antibody-stained amoebae within each ROI (**Table S2**). No amoebae were detected in ROIs from mock-infected mice (0/6, 0%) or mice examined at day 1 p.i. (0/7, 0%). At day 2 p.i., amoebae were detected in 28.6% of ROIs (2/7), increasing to 61.5% (8/13) at day 3 p.i., primarily within olfactory regions (**Fig 1C**). By day 4 p.i., amoebae were detected in 62.5% of ROIs (15/24), including regions beyond the olfactory system, such as the striatum and isocortex. Extensive hemorrhagic and necrotic damage was also observed in the olfactory bulbs of three animals examined at day 4 p.i., preventing recovery of intact olfactory bulb tissue for analysis (**Figs S1 and S3B**). Together, these observations are consistent with the progressive spread of *N. fowleri* from olfactory regions into more caudal brain regions, accompanied by extensive tissue damage at later infection time points.

### High *N. fowleri* burden is associated with innate immune and inflammatory transcriptional signatures

We sought to understand how the presence of amoebae in localized regions of the brain correlated with gene expression patterns. To this end, we compared gene expression profiles between ROIs from infected mice containing 10 or more amoebae, which we designated as regions of high amoeba burden (N = 6 tissue sections, n = 8 ROIs), and ROIs from mock-infected mice (N = 2 tissue sections, n = 5 ROIs) (**Fig 2A**). In total, we identified 129 differentially expressed genes (99 with greater expression (Log_2_-fold change > 0.58) and 30 with decreased expression (Log_2_-fold change < -0.58) genes, adj. P-value < 0.05) (**Fig2 B; Table S3)**. We observed increased expression of genes encoding for proteins associated with various families in ROIs with high amoeba burden compared to ROIs derived from mock-infected animals, including guanylate-binding proteins (GBPs) (*Gbp5*, *Gbp6*, *Gbp2*, and *Gbp3*), interferon regulatory factors (IRFs) (*Irf1*, *Irf7* and *Irf9*), and interferon-induced proteins with tetratricopeptide repeats (IFITs) (*Ifitm3*, *Ifit3*, *Ifit2* and *Ifit1*) (GBP adj. P-value = 5.66e-07; IRF adj. P-value = 1.2e-4; IFIT adj. P-value = 5.66e-07) (**Table S4**).

**FIG 2.**
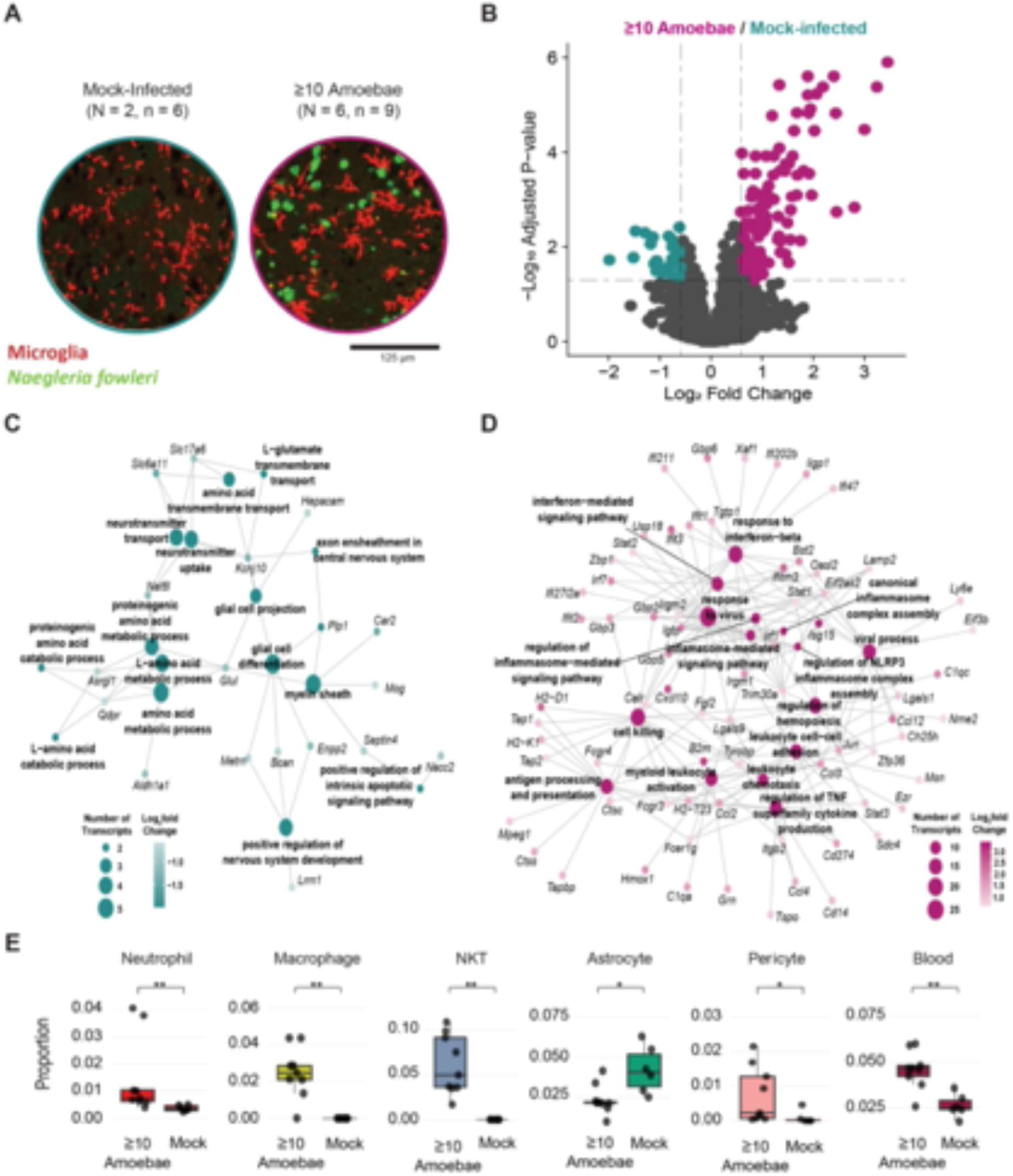
High amoeba burden is associated with pro-inflammatory transcriptional signatures. (A) Representative images of ROIs from mock-infected animals (N = 2 mouse brain tissue sections, n = 5 ROIs) and ROIs from with infected-animals with ≥ 10-amoebae (N = 6 mouse brain tissue sections, n = 8 ROIs). Colored circles highlight the representative area which was used for transcript isolation and sequencing. (B) Volcano plot displaying the differentially expressed genes between high amoeba burden ROIs (10 or more detected amoebae in the ROI) (n = 8) and ROIs from mock-infected animals (n = 5). Postive log2-fold change values (pink dots): greater expression in high amoeba burden ROIs; negative log2-fold change values (teal dots): decreased expression in high amoeba burden ROIs. (C) Network map displaying significantly depleted genes and gene set terms in high amoeba burden ROIs. The top fifteen simplified GO terms for the 30 depleted genes are shown. Genes were considered decreased if they had a less than -log2(1.5)-fold change in expression and an adjusted p-value < 0.05. (D) Network map displaying significantly enriched genes and gene set terms in high amoeba burden ROIs. The top fifteen simplified GO terms for the 99 enriched genes are shown. Genes were considered enriched if they had a greater than log2(1.5)-fold increase in expression and an adjusted p-value < 0.05. (E) Predicted proportions of specific immune and neural cell types. ROIs with high amoeba burden in brain tissue from mice ≥ 10 amoebae) were compared to ROIs from mock-infected brain tissue. Predicted cell proportions were generated using a reference dataset which combined the Immune Atlas ImmGen Cell Family and Allen Brain Atlas datasets. Statistical significance was assessed using the Wilcoxon signed-rank statistical test with BH correction.

To identify biological processes and pathways associated with high amoeba burden, we performed functional enrichment analysis. We scored 19,155 GO term gene sets against the genes differentially expressed in ROIs with 10 or more amoebae versus ROIs derived from mock-infected animals. We detected a total of 983 differentially enriched gene sets (Benjamini–Hochberg (BH) < 0.05) (**Table S5**). Gene set enrichment maps highlight distinct biological themes associated with the presence of *N. fowleri* in brain tissue (**Figs 2C and 2D**). In ROIs containing amoebae, we observed positive enrichment of gene sets associated with immune responses, including 29 terms associated with interferon signaling and its regulation, and 29 terms associated with antigen processing and presentation (**Fig 2D; Table S5**). Consistent with these findings, Reactome Pathways analysis revealed enrichment of genes involved in interferon responses and interferon stimulated signaling (**Fig S5A; Table S6**). Conversely, ROIs containing amoebae exhibited negative enrichment of gene sets associated with neurotransmitter uptake and transport (*Slc6a11*, *Kcnj10*, *Nat8l*, *Slc17a6*), as well as glial cells processes including gliogenesis, differentiation, cellular projection (*Plp1*, *Kcnj10*, *Enpp2*, *Bcan*, *Metrn, Hepacam, Glul*) (**Fig 2C**; **Table S6**). Further comparison with cell-type-specific gene signatures from the Cellmarker database identified negative enrichment of terms associated with astrocyte-associated genes (*Bcan*, *Kcnj10*, *Hepacam*, *Nacc2*, *Aldh1a1*, *Slc6a11*, *Tmem229a*, *Glul*, *Gstm5*) and two with oligodendrocyte-associated genes (*Mog*, *Plp1*, *Gpr37*, *Enpp2*, *Ank1*) (**Fig. S5D; Table S7**).

Next, we identified putative transcriptional regulatory networks associated with the differentially expressed genes in ROIs containing amoebae by searching for transcription factor (TF)-gene interactions using the Transcriptional Regulatory Relationships Unraveled by Sentence-based Text mining (TRRUST) database. We observed significant enrichment of 68 mouse TFs with potential roles in governing the gene expression changes observed in ROIs containing amoebae (adjusted p value < 0.05) (**Table S8**). The most enriched transcription factors included STAT1 (*Socs3*, *Cxcl10*, *Irf1*, *Mt1*, *Gbp2*, *Irf9*; adj. P-value = 4.43e-8) and JUN (*Socs3*, *Jun*, *Ccl12*, *Hmox1*, *Ccl2*, *Cd14*, *Gfap*; adj. P-value = 1.95e-4) (**Fig S6A; Table S8**). The regulation of gene expression through STAT1, a TF activated by IFN-y and interactor with GBP5 may drive a pro-inflammatory response(24).

Gene expression signatures provide insights into immune and neural processes associated with *N. fowleri* invasion. However, many transcripts are expressed across multiple cell types, making it difficult to distinguish their relative contributions to the observed transcriptional responses. To better characterize the cellular composition of infected brain tissue, we performed cell-type deconvolution to estimate the relative abundance of different cell populations within each ROI and compared these estimates with those from mock-infected mice. Cell-type deconvolution revealed transcriptional contributions from prominent neural cell types, including astrocytes (54/54 ROIs, 100%), excitatory neurons such as CA3 glutamatergic neurons (52/54 ROIs, 96%), *Meis2* inhibitory neurons (54/54 ROIs, 100%), and oligodendrocytes (22/54 ROIs, 40%) (**Table S9**). Additionally, this analysis predicted contributions from several immune and vascular-associated cell types across ROIs, including macrophages (29/54 ROIs, 54%), neutrophils (54/54 ROIs, 100%), natural killer T (NKT) cells (29/54 ROIs, 54%), microglia (54/54 ROIs, 100%), blood endothelial cells (53/54 ROIs, 98%), dendritic cells (38/54 ROIs, 70%) and pericytes (36/54, 67%) (**Table S9**).

We next compared estimated cell-type proportions across infection time points and amoebae burden categories. Comparing ROIs with *≥* 10 detectable amoebae to mock-infected controls revealed macrophages (adj. P-value = 5.85e-3), neutrophils (adj. P-value = 5.85e-3), dendritic cells (adj. P-value = 5.85e-3) and NKT cells (adj. P-value = 5.85e-3) as the immune cells which most strongly correlated with amoebae burden (**Fig 2E and Fig S4A; Table S10**). Increasing proportions of these predicted immune cell types also appeared to strongly correlate with increasing amoeba burden (**Fig S4B**). This analysis additionally identified trends in neural cell types, with a smaller predicted proportion of astrocytes (adj. P-value = 2.15e-2) in high amoebae burden ROIs (**Fig 2E** and **Fig S4A; Table S10**). These observations contrasted to other neural cell populations, which saw an increase in blood endothelial cells (adj. P-value = 5.85e-3) and pericytes (adj. P-value = 3.59e-2) in high amoebae burden ROIs (**Fig 2E and Fig S4A; Table S10)**. Through these analyses, we detected differences in immune cell proportions and gene expression signatures in ROIs in amoebae-infected brain tissue.

### Within-tissue analysis reveals immune-associated gene expression differences across regions with varying amoeba burden

We next investigated whether immune-associated transcriptional differences also were evident between regions of high amoeba burden and regions with no detectable amoebae within the same tissue section. Both groups of ROIs were derived from infected brain tissue, allowing us to examine spatial differences in host responses independent of infection status at the tissue level. We grouped ROIs from brain tissue sections collected on days 3 and 4 p.i. (N = 6 tissue sections) according to high amoeba burden (n = 8) or no detectable amoebae (n = 14) and analyzed them using the differential gene expression and cell-type prediction workflows described prior (**Fig 3A**). The within-section comparison identified fewer differentially expressed genes than the comparison between high amoeba burden and mock-infected ROIs, with 43 versus 99 enriched genes and 1 versus 30 depleted genes, respectively (**Fig 3B**; **Table S11**). GO term enrichment analysis revealed several terms related to immune cascades through interferon, including response to Type I interferon (*Zbp1*, *Oasl2*, *Ch25h*, *Usp18*, *Ifitm3*, *Irf7*, *Isg15*) and response to interferon-beta (*Ifi47*, *Gbp3*, *Xaf1*, *Ifi211*, *Ifi202b*, *Ifit1*, *Ifitm3*, *Bst2*, *Ifit3*) (**Fig 3C**; **Table S12**). We also observed enrichment of genes associated with eosinophil chemotaxis (*Ccl3*, *Ccl4*, *Ccl2*, *Ccl12*) (**Fig 3C**; **Table S12**), consistent with previous reports of eosinophil-associated immune responses during PAM (9). Additionally, we observed increasing expression of interferon-responsive genes (*Isg15*, *Irf7*, *Ifit3*) and chemokine genes (*Ccl3*, *Ccl4*, *Ccl2*, *Ccl12*) with increasing amoeba burden. Expression was higher in ROIs with low amoeba burden than in those without detectable amoebae and highest in ROIs with high amoeba burden (**Fig 3D**). Reactome pathway analysis identified enrichment of Type I and II interferon signaling pathways among genes associated with increasing amoeba burden, including interferon alpha/beta signaling (*Ifitm3*, *Bst2*, *Irf7*, *Isg15*, *Ifit1*) and interferon gamma signaling (*Irf7*, *B2m*, *Irf9*, *Gbp3*) (**Fig S5B**; **Table S6**). Similarly, TRRUST analysis identified JUN (adjusted *P* = 1.95e-3) and NF-*κ*B (adjusted *P* = 6.71e-3) as the most significantly enriched transcriptional regulators, suggesting their potential involvement in host transcriptional responses associated with increasing amoeba burden (**Fig S6B**; **Table S8**).

**FIG 3.**
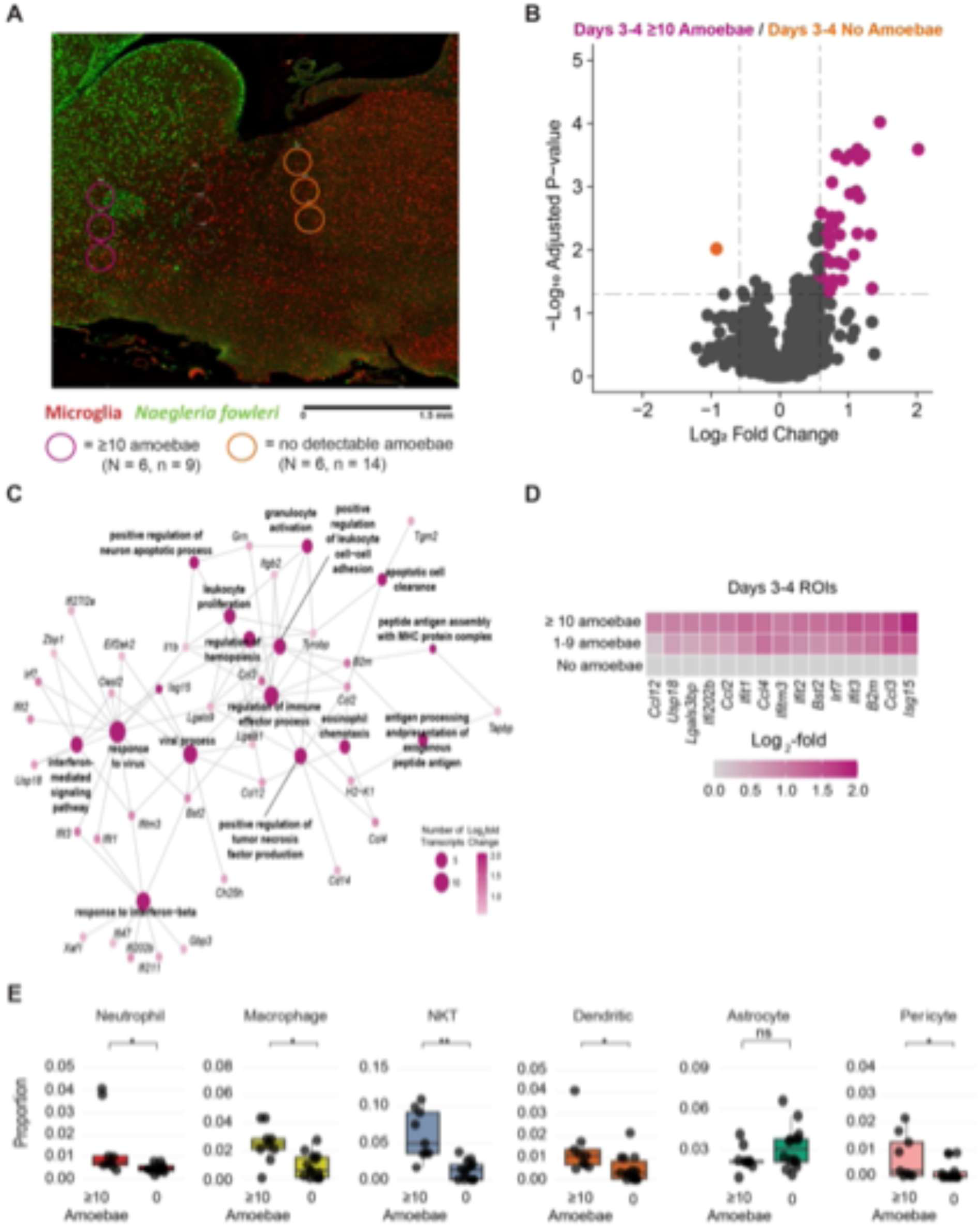
Within-tissue analysis reveals pro-inflammatory gene expression signatures associated with amoeba burden. (A) Representative image of no amoebae (n = 14 ROIs) and >10-amoebae (n = 8 ROIs) classifications in Days 3-4 mouse brain tissue sections (N = 6). Colored circles highlight the representative area which was used for transcript isolation and sequencing. (B) Volcano plot displaying differentially expressed genes between high amoeba burden ROIs (10 or more detected amoebae in the ROI) (n = 8) and no amoebae ROIs (n = 14). Postive log2-fold change values (pink dots): greater expression in high amoeba burden ROIs; negative log2-fold change values (orange dot): decreased expression in high amoeba burden ROIs. (C) Network map displaying significantly enriched genes and gene set terms in high amoeba burden ROIs. The top fifteen simplified GO terms for the 43 enriched genes are shown. Genes were considered enriched if they had a greater than 1.5 Log2-fold increase in expression and an adjusted p-value < 0.05. (D) The most enriched genes reveals a gradient of expression which correlates increasing gene expression with increasing amoeba burden. Expression was normalized to the No Amoebae ROIs within Days 3-4 mouse brain tissue sections. (E) Predicted proportions of select responsive immune cells and neural cells. ROIs with high amoeba burden in brain tissue from mice (>10 detectable amoebae) were compared to ROIs with no amoeba within the same brain tissue section. Predicted cell proportions were generated using a reference dataset which combined the Immune Atlas ImmGen Cell Family and AllenBrainAtlas datasets. Statistical significance was assessed using the Wilcoxon signed-rank statistical test with BH correction.

Cell-type deconvolution revealed immune-cell profiles similar to those observed when comparing ROIs with high amoeba burden to mock-infected controls (**Fig 3E and Fig S7A**). Specifically, ROIs with high amoeba burden had significantly higher predicted proportions of macrophages (adjusted *P* = 3.17e-2), neutrophils (adjusted *P* = 3.17e-2), dendritic cells (adjusted *P* = 3.13e-2), pericytes (adjusted *P* = 3.29e-2), and NKT cells (adjusted *P* = 9.52e-2) than ROIs without detectable amoebae from the same tissue sections (**Fig 3E and Fig S7A**; **Table S10**). Conversely, ROIs with high amoeba burden showed a modest, nonsignificant reduction in the predicted proportion of astrocytes (adjusted *P* = 3.38e-1). Together, these analyses revealed spatial differences in immune-associated transcriptional signatures within infected brain tissue, with greater inflammatory gene expression in regions containing amoebae. However, these differences were less pronounced between ROIs with and without detectable amoebae within infected brains than between high-burden ROIs and mock-infected controls. These findings suggest that inflammatory responses extend beyond regions of detectable amoeba invasion and may develop before amoebae reach those areas.

### Early immune-associated transcriptional signatures extend beyond regions of detectable amoeba invasion

Our preceding analyses focused on gene expression and inferred cell-type composition in brain regions with high amoeba burden, which may represent sites of active invasion, inflammation, and tissue damage. To determine whether host immune responses extend beyond these regions, we next examined immune-associated transcriptional signatures in areas of infected brain tissue without detectable amoebae.

We first assessed temporal changes in ROIs without detectable amoebae by grouping samples collected at days 1–2 p.i. (N = 4 tissue sections; n = 11 ROIs) and days 3–4 p.i. (N = 6 tissue sections; n = 14 ROIs) for differential gene expression and cell-type deconvolution analyses (**Fig 4A**). We identified 102 differentially expressed genes, including 85 with increased expression (log_2_ fold change > 1.5) and 17 with decreased expression (log_2_ fold change < -1.5) at days 3–4 p.i. (adjusted *P* < 0.05) (**Fig 4B**; **Table S13**). Gene Ontology (GO) enrichment analysis identified terms associated with response to interferon-beta (*Gbp2*, *Igtp*, *Ifitm3*, *Irgm1*, *Gbp6*, *Ifit3*, *Iigp1*, *Irf1*, *Gbp3*, *Tgtp1*) and response to Type II interferon (*Gbp2*, *Igtp*, *Ifitm3*, *Gbp5*, *Irgm1*, *Gbp6*, *Irf1*, *Gbp3*, *Vim*, *Ccl2*, *Zyx*, *Tgtp1*), highlighting both Type I and II interferon-associated transcriptional responses (**Fig 4D**; **Table S14**). We also identified ten enriched GO terms associated with inflammasome-mediated signaling (*Hspa8*, *Irgm1*, *Gbp5*, *Igtp*, *Gbp2*) (**Table S14**). Examination of gene expression across individual time points revealed increased expression of these immune-associated genes beginning at day 3 p.i., suggesting that inflammatory transcriptional responses develop in regions without detectable amoebae as infection progresses (**Fig 4E**). Conversely, negatively enriched GO terms were associated with neuronal dendritic processes, including regulation of dendrite extension (*Cpne6*, *Syt17*; 3 terms) and the dendritic shaft (*Jph4*, *Grm5*) (**Fig 4C**; **Table S14**). Consistent with these findings, expression of the associated transcripts decreased at days 3–4 p.i. (**Fig 4E**).

**FIG 4.**
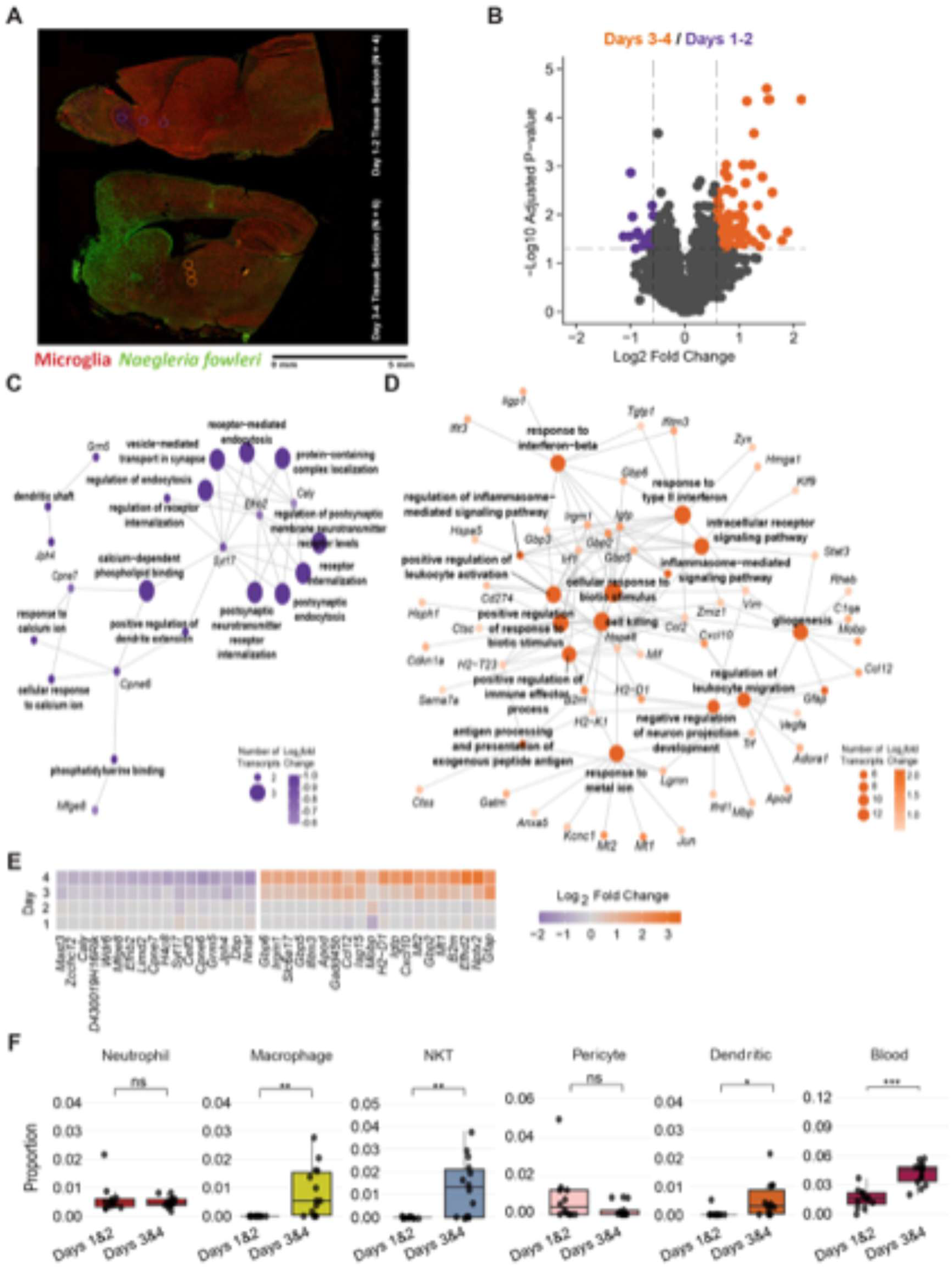
Early-stage immune-response related gene expression signatures are detectable in regions of infected brain with no amoeba. (A) Representative images of early–stage (Days 1-2, N = 4 mouse brain tissue sections, n = 11 ROIs) and late-stage (Days 3-4, N = 6, n = 14 ROIs) no amoebae classifications in mouse brain tissue sections (N = 6). Colored circles highlight the representative area which was used for transcript isolation and sequencing. (B) Volcano plot of differentially expressed genes between late-stage (Days 3-4, n = 14) and early stage (Days 1-2, n = 11) ROIs from infected animals that do not have detectable amoebae burdens. Postive log2-fold change values (orange dots): greater expression in days 3-4 ROIs; negative log2-fold change values (purple dots): decreased expression in days 3-4 ROIs. (C) Network map displaying significantly genes and gene set terms in Days 3-4 no amoeba burden ROIs. The top fifteen simplified GO terms for the 17 depleted genes are shown. (D) Network map displaying significantly enriched genes and gene set terms in Days 3-4 no amoeba burden ROIs. The top fifteen simplified GO terms for the 85 enriched genes are shown. Genes were considered enriched if they had a greater than 1.5 Log2-fold increase in expression and an adjusted p-value < 0.05. Genes were considered depleted if they had a less than -1.5 Log2-fold increase in expression and an adjusted p-value < 0.05. (E) The most enriched genes reveals a gradient of expression which correlates with increasing gene expression over time, whereas the most depleted genes reveals a gradient of expression which correlates with decreasing gene expression over time. The expression of a gene on each day was normalized to the average expression of that gene across ROIs collected on Days 1-2. (F) Predicted proportions of immune cells and neural cells. ROIs with no amoeba burden in brain tissue from mice euthanized on Days 3-4 were compared to ROIs with no amoebae from mice euthanized on Days 1-2. Predicted cell proportions were generated using a reference dataset which combined the Immune Atlas ImmGen Cell Family and AllenBrainAtlas datasets. Statistical significance was assessed using the Wilcoxon signed-rank statistical test with BH correction.

Reactome pathway analysis revealed enrichment patterns similar to those identified when comparing ROIs with high amoeba burden to mock-infected controls (**Fig S5C**; **Table S6**). We also identified pathways associated with cellular stress responses, including Cellular Responses to Stress (*Hspa8*, *Cdkn1a*, *Jun*, *Hspa5*, *Stat3*, *Hmga1*, *Higd1a*, *Hsp90b1*, *Vegfa*, *Hsph1*, *Rheb*, *Ccl2*, *Igfbp7*), activating transcription factor 6 (ATF6)-associated responses (*Hspa5*, *Hsp90b1*; 2 terms), creatine metabolism (*Gatm*, *Slc6a8*), and regulation of gene expression by hypoxia-inducible factors (*Higd1a*, *Vegfa*) (**Fig S5C**; **Table S6**). Complementary analysis using the TRRUST database identified NFIC (*Ccl2*, *Mbp*, *Mt1*, *Gfap*; adjusted *P* = 2.22e-5) and NFKB1 (*Cxcl10*, *Cdkn1a*, *Ccl12*, *Gadd45b*, *Fth1*, *Adora1*, *Ccl2*, *Mbp*, *Vegfa*; adjusted *P* = 2.22e-5) as the most significantly enriched candidate transcriptional regulators (**Fig S6C**; **Table S8**).

We next examined whether these transcriptional changes coincided with differences in predicted cell-type composition across infection time points. Among ROIs without detectable amoebae, the predicted proportions of blood endothelial cells (*R*^2^ = 0.46, adjusted *P* = 2.7e-4), macrophages (*R*^2^ = 0.38, adjusted *P* = 7.6e-4), microglia (*R*^2^ = 0.23, adjusted *P* = 1.2e-2), and NKT cells (*R*^2^ = 0.39, adjusted *P* = 7.6e-4) increased with infection day (**Fig S8B**). When comparing early (days 1–2 p.i.) and later (days 3–4 p.i.) infection time points, we identified significantly higher predicted proportions of macrophages (adjusted *P* = 3.82e-3), dendritic cells (adjusted *P* = 1.95e-2), and NKT cells (adjusted *P* = 6.35e-3) in ROIs collected at later time points (**Fig 4F**; **Fig S8A**; **Table S10**). In contrast, predicted neutrophil proportions did not differ significantly between these groups (adjusted *P* = 8.6e-1), suggesting potential differences in the temporal patterns of immune-cell-associated responses during PAM progression. Notably, blood endothelial cells showed a highly significant increase in predicted proportions at later infection time points (adjusted *P* = 1.49e-4) (**Fig 4F and Fig S8A**; **Table S10**). Together, these findings suggest that endothelial- and immune-cell-associated transcriptional responses develop in brain regions without detectable amoebae as PAM progresses, potentially preceding localized amoeba invasion.

## DISCUSSION

In this study, we used spatial transcriptomics to characterize host gene expression across brain regions and over the course of PAM progression in mice. We identified transcriptional signatures associated with developing pro-inflammatory responses to *N. fowleri* infection and, through cell-type deconvolution, inferred the relative contributions of immune cell populations across tissue regions with varying amoeba burdens. Notably, we found that immune-associated transcriptional signatures were evident in regions of infected brain tissue without detectable amoebae, suggesting that host responses extend beyond areas where amoeba exist.

PAM presents a challenge for the host immune system, requiring the elimination of an invasive pathogen while limiting damage to the brain. Previous studies have implicated several components of the immune response to *N. fowleri*, including neutrophils (12), macrophages (13), microglia (15), and the complement system (25, 26). Consistent with these findings, our cell-type deconvolution analysis suggested that the inferred contributions of macrophages and neutrophils varied with local amoeba burden, while also identifying a potential contribution from NKT cells. These findings suggest that a diverse array of immune cell populations may contribute to the host response during *N. fowleri* infection. Notably, endothelial-associated transcriptional signatures were evident early during infection, including in brain regions without detectable amoebae. Macrophage-, NKT cell-, and dendritic cell-associated signatures were also identified in regions lacking detectable amoebae, suggesting that immune-associated responses extend beyond areas of direct invasion. Although the factors underlying these spatially distributed responses remain unclear, previous *in vitro* studies have demonstrated that host cells can respond to factors secreted by *N. fowleri* (27, 28). Alternatively, signals released from damaged neural tissue may contribute to local inflammatory responses (29). The reduced inferred contribution of astrocytes in amoeba-rich regions may reflect accompanying changes in tissue composition or cellular transcriptional states. Together, these findings suggest that immune-associated transcriptional responses extend beyond regions containing amoebae, with endothelial-associated signatures potentially reflecting an early component of this spatially distributed host response.

Among the cell populations identified through deconvolution, the inferred contribution of blood endothelial cells increased over time in regions without amoebae and was higher in ROIs with high amoeba burden than in mock-infected tissue. These findings suggest that vascular-associated transcriptional changes are evident during early *N. fowleri* infection, including in regions without amoebae. Although the blood endothelial reference profile in the ImmuneAtlas ImmGen Cell Family dataset represents a broad vascular signature rather than a blood-brain barrier-specific endothelial population, these observations highlight a potential association between vascular-associated transcriptional responses and the spatial progression of infection. Similar changes in macrophage-, NKT cell-, and other immune cell-associated signatures further suggest that the host response involves multiple cellular populations, including those associated with both resident brain tissue and peripheral immunity.

Interferon-associated transcriptional signatures were prominent in infected mouse brain tissue during later stages of disease progression, including regions with differing amoeba burdens. Previous studies have implicated IFN-*γ* in the host response to *N. fowleri* (3, 16). Additionally GO enrichment analyses identified pathways associated with type I interferon signaling, including responses to IFN-*α* and IFN-*β*. Consistent with these findings, we observed increased expression of several interferon-associated gene families, including interferon regulatory factors (IRFs) (30), guanylate-binding proteins (GBPs) (24), and interferon-induced proteins with tetratricopeptide repeats (IFITs) (31). Increased expression of IRF and IFIT family members has also been reported in previous transcriptomic analyses of *N. fowleri*-infected brain tissue (16), further supporting the prominence of these transcriptional signatures during infection. Interestingly, type I interferons have been implicated in the regulation of NKT cell responses (32), providing a potential connection to the NKT cell-associated signatures identified through our deconvolution analysis. Together, these findings highlight interferon-associated transcriptional programs as a prominent feature of the host response to *N. fowleri*, extending across brain regions with varying degrees of amoeba invasion.

The enrichment of type I interferon-associated transcriptional signatures during *N. fowleri* infection is notable, as these pathways traditionally are associated with antiviral immunity, but may contribute to host responses against nonviral pathogens (33). Similar pro-inflammatory transcriptional signatures have been reported in other parasitic diseases affecting the central nervous system, including *Toxoplasma* encephalitis and cerebral malaria (34, 35). While previous studies of these conditions have highlighted contributions from adaptive immune populations, including CD8^+^ T cells and B cells (36, 37), our findings predominantly revealed transcriptional signatures associated with innate immune responses. Of particular relevance, a spatial transcriptomic study of cerebral malaria identified endothelial-associated inflammatory signatures involving chemokine expression and antigen presentation, with potential implications for cerebral edema (35). Similarly, our analysis revealed increased inferred contributions from vascular endothelial cells, elevated expression of *Ccl3* and *Ccl4* transcripts, and enrichment of GO terms related to antigen presentation. These parallels suggest that vascular-associated inflammatory responses may be a shared feature of parasitic diseases affecting the brain and could be relevant to the pronounced cerebral edema observed in PAM (11).

Despite the prominent immune-associated transcriptional signatures observed during *N. fowleri* infection, all mice examined at day 4 p.i. had reached end-stage disease criteria, indicating that these responses did not prevent severe disease in this experimental model. The characteristics of an effective host response to *N. fowleri* remain poorly understood, particularly whether the timing or magnitude of inflammatory responses influences disease progression. Previous studies have implicated IFN-*γ*-associated inflammatory responses during infection (16, 8). Although these responses may contribute to host defense, excessive inflammation within the brain could also exacerbate tissue damage and neurological dysfunction. Our findings reveal extensive inflammatory transcriptional signatures during advanced infection but do not establish whether these responses contribute to parasite control, disease pathology, or both.

Differences in host genetic background and parasite virulence may provide additional insight into the relationship between immune responses and disease outcomes. Previous studies have reported differences in survival among mouse strains infected with the same *N. fowleri* isolate, with CBA/J and C57BL/6J mice surviving longer than FVB/NJ and NOD/ShiLtJ mice (38). These findings suggest that host genetic background may influence susceptibility to infection, potentially through differences in immune responses or other host factors. Similarly, *N. fowleri* isolates exhibit considerable variation in virulence. The V596 isolate used in this study is highly virulent, with reported disease progression occurring within approximately 4–5 days in CD-1/ICR mice, whereas infection with the less virulent V067 isolate has been associated with survival durations of approximately 10–12 days (39). These differences raise the possibility that the magnitude, timing, or composition of host inflammatory responses may vary with parasite virulence and contribute to differences in disease progression. However, whether the pronounced inflammatory signatures observed following V596 infection reflect responses to greater parasite-mediated tissue damage or contribute independently to disease severity remains unresolved.

Together, these findings provide a foundation for further investigations into the relationship between host inflammatory responses and disease progression during PAM. These new spatial transcriptomic data represent a resource for examining regional and temporal differences in host gene expression, while future studies employing single-cell approaches could help distinguish changes in cellular abundance from altered transcriptional states and identify the cellular sources of immune-associated transcripts. Building on the limited transcriptomic studies of *N. fowleri* across different isolates and parasite stages (40, 16, 41, 42), future spatial ‘omic approaches incorporating both host- and amoeba-derived signals could enable the identification of coordinated molecular signatures within infected tissues. Such analyses, extended to genetically diverse hosts and *N. fowleri* isolates with differing virulence, could further clarify how parasite-associated factors and the timing and magnitude of host inflammatory responses relate to disease outcomes. Ultimately, defining the contributions of host inflammatory responses to parasite control and disease pathology may help identify new opportunities for host-directed interventions against PAM.

## MATERIALS AND METHODS

### *N. fowleri* isolates and cultures

We used the *N. fowleri* V596 isolate (Genotype III), obtained from a male patient who succumbed to PAM in 2007 in Nevada, USA. This isolate causes an end-stage infection in 4-6 days in CD-1 (ICR) mice (39). The isolate was provided by Dr. Ibne Ali (Centers for Disease Control, Atlanta, GA, USA). We maintained parasites axenically in Nelson’s complete media (NCM) supplemented with 10% Fetal Bovine Serum (FBS) and 1% penicillin-streptomycin as described previously (43).

### Animal husbandry and infection procedures

We conducted animal studies in the laboratory animal facilities at the University of Georgia’s College of Veterinary Medicine (CVM). The studies were approved by the Institutional Animal Care and Use Committee (IACUC) (A2024-03-003-Y3-A2). We housed mice in conventional cages, 22.2°C +/- 2°C and 50% +/- 20% relative humidity with a controlled 12-h light / 12-h dark cycle.

For infection of mice, we isolated amoebae from axenic culture by icing the flask for 10 minutes, pelleted via centrifugation at 3900 RPM for 5 minutes at room temperature, and washed 2 times in phosphate buffered saline (PBS) prior to seeding onto a flask containing a confluent monolayer of Vero cells (green monkey kidney cells; E6; ATCC CRL-1586). We cultured Vero cells as previously described (44). We monitored the amoeba-Vero cell co-cultures daily and sub-cultured them once >85% of the Vero cells had been destroyed by the amoebae. To “prime” amoebae for infection, we serially passaged amoebae 6 times over Vero cells (39). Upon completion of the sixth passage, we collected amoebae via centrifugation, re-suspended the pellet containing amoebae in 500 µL of Dulbecco’s Modified Eagle Medium (DMEM). When necessary, we adjusted the concentration of the suspension to 500 amoebae per µL. We inoculated 10 µL of the parasite suspension into a single nare of an anesthetized female 3-4-week-old ICR (CD-1) mouse (n = 10). As controls, we mock-infected two mice with DMEM alone. We monitored animals’ mass daily and performed inspections (three times per day) for symptoms of PAM, including weight loss, piloerection, orbital tightening, arched spine, ataxia, increased respiratory effort and seizures or other neurological symptoms as described previously (45).

### Tissue collection and slide generation

We sacrificed mice at time-points 24-hour intervals post-inoculation over the 4-day course of infection. We sacrificed one mouse at random on Days 1 and 2, and three mice randomly on Day 3, as no mice were displaying symptomatic criteria. On the fourth day p.i., we euthanized three mice for displaying symptoms of PAM (per the approved animal use protocol). Immediately upon sacrifice, we transcardially perfused mice with 50 mL of PBS using a syringe affixed with a 27-gauge needle. We then dissected the brain and placed it into 10 mL of 10% neutral-buffered formalin (NBF; pH 7.0) solution for 24-48 hours at room temperature. We processed mock-infected brains alongside mice displaying end-stage symptomology on day 4 post-infection (p.i.). After formalin fixation, we further processed brains by sagittal and frontal dissection to isolate tissue from the infected lobe. Processed tissue was then embedded into formalin-fixed, paraffin-embedded (FFPE) tissue blocks, and five µm sections were cut and adhered to Leica® BOND plus slides.

### Preparation of brain tissue for transcriptomic spatial profiling

We submitted slides containing brain tissue to the Bruker (formerly Nanostring) Customer Experience Lab (Seattle, WA) for processing through the GeoMx Digital Spatial Profiler (DSP) platform. Slides were dried overnight, and the next day incubated at 60 °C for 30 minutes, dewaxed using Citrisolv, dried in 100% and 95% ethanol washes, and then washed once with PBS. For antigen retrieval, slides were briefly submerged in 99 °C DEPC-H2O, then placed in a 99°C 1X Tris-EDTA (pH 9.0) solution in a steamer for 15 minutes. The samples were then subjected to enzymatic exposure of RNA targets for 15 minutes at 37 °C in a 1 *µ*g/mL Proteinase K solution. Samples were then immediately fixed with 10% NBF for 5 minutes, then processed through two five-minute washes in a tris-glycine solution (stop buffer) and a PBS wash. GeoMx WTA RNA probes (531–121401119) were added to the slides before an overnight incubation in a 37 °C hybridization oven. The following morning, the slides went through two rounds of a 2x Saline Sodium Citrate (SSC)/50% formamide stringent wash followed by two brief washes in 2X SSC. Blocking was conducted with GeoMx Buffer W for 30 minutes.Three antibodies were used to label amoebae and tissue architecture of the brains: a *N. fowleri*-specific antibody conjugated to an AF532 fluorophore to detect amoebae; an AF647 TMEM-119 antibody (Synaptic Systems; 400 011) to detect microglia; and an AF594 Ly-6G antibody (Biolegend; 127607) to stain neutrophils. These markers and a DNA stain (Syto13, 531-121300303) were then incubated on the slides for 60 minutes followed by two washes in 2X SSC. The slides were loaded onto a GeoMx Digital Spatial Profiler for region of interest (ROI) selection and data acquisition. Once the samples were imaged and regions of interest were confirmed, each ROI was exposed to a UV LED light, which releases the barcoded oligos for capture by microfluidics. The UV LED light was collimated to be reflected from the digital micromirror device and focused at the sample tissue for 1-10 seconds at a 365-385 nm wavelength (46). The released barcodes were then collected into individual wells, by ROI, in a 96-well DSP collection plate in preparation for library processing and sequencing.

### Library preparation and sequencing

DSP collection plates were dried down and then each collection well was resuspended in 10*µ*L of water. A library was generated using 4*µ*L from each of the DSP collection wells and purified via a magnetic bead-based protocol. The resulting library was then sequenced on a NextSeq 2000 using reversed 2×27 base paired end reads. After sequencing, the resultant FASTQ files were converted into Digital Code Counts (DCCs) and uploaded to the GeoMx instrument for analysis via the Digital Spatial Profiler Data Analysis software (Version 3.1.2.12) on-instrument.

### ROI image processing and data normalization

We loaded three sets of data into R (Version 4.4.1) for analysis of transcripts isolated from the placed ROIs. This data includes an ImageJ spreadsheet detailing the numbers and size of amoebae counted within the ROIs (**Table S2**). Briefly, we downloaded images of each ROI from the GeoMX DSP Analysis Suite. First, we overlayed the ROI multichannel composite images into one image. We then selected amoebae in the appropriate channel, and a threshold was created to eliminate background staining. We then assigned each ROI to a brain region or structure according to the labelled map found in **Fig S1B**. An example image was manually annotated using the Allen Institute Brain Atlas sectioning map (Mouse, P56, Sagittal) as a reference (**Fig S3A**). A schematic of the ImageJ pipeline we used to count amoebae can be found in **Figure S2**. We then downloaded the normalized transcript reads using the Bruker GeoMx Digital Spatial Profiler (DSP; Version 3.1.2.12).

For transcriptomic analyses, we exported gene expression data from the Bruker GeoMx Digital Spatial Profiler (DSP; version 3.1.2.12) and normalized the data according to the parameters described in **Table S15** in the supplemental material. The unnormalized and third-quartile (Q3)-normalized expression data are provided in **Tables S16 and S17** in the supplemental material, respectively. We then imported the transcriptomic data alongside the ROI-associated image measurements into R (version 4.4.1) for downstream analyses.

### Differential gene expression analyses

We classified ROIs according to sampling day and amoeba burden to define groups for differential gene expression analyses. Amoeba burden was categorized as zero (0), low (1-9 amoebae), or high (10 or more amoebae). ROIs from uninfected control mice were assigned an amoeba burden of zero. Using the GeoMx DSP software (version 3.1.2.12), we performed differential gene expression analyses across three comparisons: (i) Days 3–4 ROIs with high amoeba burden versus uninfected control ROIs (**Table S2A**); (ii) Days 3–4 ROIs with high amoeba burden versus Days 3–4 ROIs with no detectable amoebae (**Table S2B**); and (iii) Days 3 and 4 ROIs with no detectable amoebae versus Days 1 and 2 ROIs with no detectable amoebae (**Table S2C**). We defined differentially expressed genes as those with an absolute log2 fold change greater than log_2_(1.5) or less than -log_2_(1.5) and a BH-adjusted *P* value *<* 0.05.

### Functional enrichment analyses

We performed Gene Ontology (GO) enrichment analyses using clusterProfiler (version 4.14.6) (47) and the organism-specific mouse annotation database org.Mm.eg.db (version 3.20.0) (48). We additionally used Reactome Pathways (2024) (49), CellMarker (2024) (50), and TRRUST Transcription Factors (2019) (51) to investigate biological pathways, cellular signatures, and transcriptional regulatory processes associated with differential gene expression across conditions. We used enrichment maps (52) to visualize functionally related gene sets and identify broader biological themes. We used We applied the Benjamini-Hochberg (BH) method (53) to correct for multiple testing and considered enrichment results with an adjusted *P* value < 0.05 statistically significant.

### Cell-type deconvolution and statistical analyses

To estimate cell-type proportions within individual ROIs, we performed spatial deconvolution using the NanoString SpatialDecon package (version 1.16.0). We used two reference datasets available through the NanoString-Biostats/SpatialDecon library: the ImmuneAtlas ImmGen Cell Family dataset, which contains immune cell gene expression profiles from the Immunological Genome Project (54), and the AllenBrainAtlas dataset, which contains single-cell gene expression profiles of neural cell populations annotated by the Allen Institute for Brain Science (55). We combined these datasets into a single reference matrix to enable the simultaneous inference of immune and neural cell populations (**Table S18**).

We used the Wilcoxon signed-rank test with BH correction (53) to evaluate differences in predicted immune cell compositions between ROIs. For simple linear regression analyses, we used F-tests with BH correction to assess statistical significance. Results of all statistical tests are provided in **Table S6** in the supplemental material.

## ETHICS APPROVAL

We conducted animal studies in the laboratory animal facilities at the University of Georgia’s College of Veterinary Medicine (CVM). The studies were approved by the Institutional Animal Care and Use Committee (IACUC) (A2024-03-003-Y3-A2).

## ACKNOWLEDGMENTS

We thank Dr. Ibne Ali (Centers for Disease Control and Prevention) for providing the *N. fowleri* V596 isolate and Dr. Ashley Moseman (Duke University) for providing the *N. fowleri-*specific antibody used to visualize amoebae in this study. We also thank the Customer Experience Lab at Bruker Spatial Biology (Seattle, WA) for technical assistance with tissue slide preparation, GeoMx Digital Spatial Profiler processing, and sequencing.

## AUTHOR CONTRIBUTIONS

Conceptualization: Joseph Dainis, Anthony A. Ruberto, Dennis E. Kyle; Data curation: Joseph Dainis, Anthony A. Ruberto; Formal analysis: Joseph Dainis, Anthony A. Ruberto; Funding acquisition: Joseph Dainis, Dennis E. Kyle; Investigation: Joseph Dainis, Anthony A. Ruberto; Methodology: Joseph Dainis, Anthony A. Ruberto, Michael Bailey, Dennis E. Kyle; Project administration: Anthony A. Ruberto, Dennis E. Kyle; Supervision: Anthony A. Ruberto, Dennis E. Kyle; Visualization: Joseph Dainis, Anthony A. Ruberto; Writing – original draft: Joseph Dainis, Anthony A. Ruberto; Writing – review & editing: Joseph Dainis, Anthony A. Ruberto, Mackenzie A. C. Sievert, Michael Bailey, Dennis E. Kyle.

## Supplemental material

**FIG S1** GeoMx imaging of mock-infected and *N. fowleri*-infected mouse brain tissue.

**FIG S2** ImageJ workflow for quantification of amoeba burden within ROIs.

**FIG S3** Anatomical localization of ROIs and gross brain pathology during *N. fowleri* infection.

**FIG S4** Predicted immune and neural cell proportions in relation to *N. fowleri* burden in the murine brain.

**FIG S5** Reactome pathway and CellMarker gene signature enrichment across regions of differing amoeba burden.

**FIG S6** TRRUST transcription factor target enrichment across ROIs differing in amoeba burden and infection time point.

**FIG S7** Predicted immune and neural cell proportions in regions with differing amoeba burdens within infected brain tissue.

**FIG S8** Predicted immune and neural cell proportions in brain regions without detectable amoebae across infection time points.

**TABLE S1** ROI Sequencing Outputs.

**TABLE S2** Amoebae Counts in ROIs Quantified by ImageJ Analysis.

**TABLE S3** Differential Gene Expression for *≥* 10 Amoebae vs Uninfected ROIs from All Mouse Brain Tissues.

**TABLE S4** Gene Family Enrichment Test Across Three Comparisons (*≥* 10 amoebae versus uninfected; *≥* 10 amoebae Days 3-4 vs No amoebae Days 3-4; No Amoebae Days 3-4 vs No Amoebae Days 1-2) using Fisher’s Statistical Test.

**TABLE S5** GO Term Enrichment for *≥* 10 Amoebae ROIs compared to Uninfected ROIs.

**TABLE S6** Reactome (2024) Term Enrichment for Three ROI Comparisons.

**TABLE S7** CellMarker (2024) Term Enrichment for Uninfected ROIs (as compared to *≥* 10 amoebae ROIs).

**TABLE S8** Transcriptional Regulatory Relationships Unraveled by Sentence-based Text mining (2019) Term Enrichment for Three ROI Comparisons.

**TABLE S9** Estimated cell type proportions for all 54 analyzed ROIs.

**TABLE S10** Wilcoxon Statistical Results for Predicted Immune Cell Proportion Changes Across Three Comparisons.

**TABLE S11** Differential Gene Expression for *≥* 10 Amoebae vs No Amoebae ROIs from Days 3-4 Infected Mouse Brain Tissue.

**TABLE S12** GO Term Enrichment for *≥* 10 Amoebae ROIs from Days 3-4 only compared to No Amoebae ROIs from Days 3-4.

**TABLE S13** Differential Gene Expression for No Amoebae ROIs in Days 3-4 versus Days 1-2 Mouse Brain Tissues.

**TABLE S14** GO Term Enrichment for No Amoebae ROIs from Days 3-4 only compared to No Amoebae ROIs from Days 1-2.

**TABLE S15** Digital Spatial Profiler Quality Control and Normalization Pipeline.

**TABLE S16** Non-normalized spatial transcriptomic data.

**TABLE S17** Q3-normalized spatial transcriptomic data.

**TABLE S18** Reference dataset details using combined Allen Brain Institute AllenBrainAtlas and ImmuneAtlas Immgen Cell Family Datasets.

## DATA AVAILABILITY STATEMENT

Raw sequencing data (FASTQ files) generated from GeoMx Digital Spatial Profiler analyses of *N. fowleri*-infected and mock-infected mouse brain tissue have been deposited in the National Center for Biotechnology Information (NCBI) Sequence Read Archive (SRA) under BioProject accession number PRJNA1559718. Processed gene expression data and associated ROI metadata are provided in the supplementary material.

